# Combined elevated temperature and CO_2_ alter *Wolbachia* abundance and host fitness but not cytoplasmic incompatibility in a stored-product pest

**DOI:** 10.64898/2026.11.01.763417

**Authors:** Phyu Phyu San, Ryosuke Fujita, Julia Ferrari, Midori Tuda

**Author notes:** Correspondence: M. Tuda, Laboratory of Insect Natural Enemies, Faculty of Agriculture, Kyushu University, Fukuoka 819-0395, Japan.

## Abstract

Coinfecting endosymbiont strains may respond differently to environmental change, with consequences for host performance and reproductive incompatibility. We investigated an isofemale line of *Callosobruchus chinensis* coinfected with the cytoplasmic incompatibility (CI)-inducing *Wolbachia* strains wBruCon (Con) and wBruOri (Ori), together with derived Con-only and uninfected sublines. Beetles were maintained under control conditions (30°C, ∼500 ppm CO_2_) or under combined elevated temperature and CO_2_ (32°C, ∼1000 ppm CO_2_; eT&CO_2_). In coinfected adults, Ori relative abundance declined under eT&CO_2_, whereas Con abundance increased in females. Con responses also depended on host sex and infection status, with singly infected males showing particularly high abundance under control conditions. However, CI expression remained stable across environmental conditions. Coinfected females laid fewer eggs than uninfected females, whereas host life-history responses to eT&CO_2_ varied among traits and generations. eT&CO_2_ accelerated development and reduced adult longevity, while reduced egg– adult survival emerged in the F2 generation. It also reduced F1 fecundity but only in the uninfected subline. Strong and partial CI associated with Con and Ori, respectively, remained evident under both environmental conditions. Equilibrium analyses incorporating relative fecundity, CI intensity, and maternal transmission suggested lower infection persistence under eT&CO_2_ in the F1 generation but higher persistence in the F2 generation. Sensitivity analysis suggested contribution of improved relative fecundity of infected females in the latter. Overall, the two *Wolbachia* strains exhibited contrasting, sex-dependent abundance responses to eT&CO_2_, while CI expression remained stable. These findings suggest that climate change may alter symbiont infection dynamics in economically important pests, with potential implications for symbiont-mediated traits, including pesticide resistance.

## 1. Introduction

*Wolbachia* are maternally inherited intracellular bacteria that infect a large fraction of terrestrial arthropods, including many beetles (Weinert et al., 2015; Kajtoch & Kotásková, 2018). Their effects on hosts range from beneficial to costly and may include manipulation of host reproduction (Werren et al., 2008). One of the best-known phenotypes is cytoplasmic incompatibility (CI), in which embryos produced by crosses between infected males and females lacking a compatible *Wolbachia* infection suffer elevated mortality (Zhang et al., 2010; Takano et al., 2011; Turelli & Hoffmann, 1995; Atyame et al., 2014). Because *Wolbachia* are transmitted through females, their persistence depends on the balance among reproductive manipulation, maternal transmission, and effects on host fitness.

The phenotypic effects of *Wolbachia* vary with host genotype, symbiont strain, and environmental conditions. Reported costs include reductions in fecundity, survival, longevity, reproduction, and body size (Fleury, 2000; de Almeida et al., 2011; Sumida et al., 2017; Tuda et al., 2021), whereas benefits include increased hatchability, fecundity, and resistance to chemical stress (Dobson et al., 2004; Dean, 2006; Miksanek & Tuda, 2023). For CI-inducing strains, infection dynamics are shaped by CI strength, maternal transmission fidelity, and the relative fitness of infected females (Jaenike, 2009; Turelli, 1994). Consequently, environmental changes that alter any of these components may alter infection persistence (Kriesner et al., 2016).

Coinfection adds further complexity because co-occurring strains may facilitate, compete with, or regulate one another. Coinfecting strains can differ in their ecological requirements and within-host distributions, leading to strain-specific responses to environmental conditions. For example, naturally coinfecting *Wolbachia* strains in *Drosophila simulans* exhibit pronounced sex-, tissue-, and stage-specific variation in abundance (Jones et al., 2022). Such differences suggest that environmental change may alter the relative success of coinfecting strains, with consequences for host fitness and symbiont persistence. Although environmental effects on *Wolbachia* have been studied extensively in aphids and *Drosophila* (Oliver et al., 2006; McLean et al., 2018; Smee et al., 2021; Jones et al., 2022), comparable studies in beetles remain scarce despite their high diversity (Kajtoch & Kotásková 2018; Ślipiński et al., 2011). In *Chelymorpha alternans*, coinfection strengthens CI relative to single infection (Keller et al., 2004), while in *Callosobruchus chinensis*, coinfection can alter *Wolbachia* density and infection may reduce fecundity in some host backgrounds (Okayama et al., 2016; Kondo et al., 2005). However, it remains unclear how environmental change affects coinfecting strains and whether such responses translate into altered host fitness or infection persistence.

Temperature is a major determinant of *Wolbachia*–host interactions (Charlesworth et al., 2019; Truitt et al., 2019). Elevated temperatures frequently reduce *Wolbachia* density or expression (Hu et al., 2023; van Opijnen & Breeuwer 1999; López-Madrigal & Duarte 2019), potentially influencing transmission fidelity, pathogenicity, and reproductive phenotypes (Narita et al., 2007; Mancini et al., 2021; Jaenike, 2009). In contrast, the effects of elevated atmospheric CO_2_, alone or in combination with temperature, remain poorly understood. Evidence from the linyphiid spider *Hylyphantes graminicola* suggests that elevated CO_2_ can interact with *Wolbachia* infection to alter host physiology (Su et al., 2019), indicating that ongoing climate change may influence symbiont–host interactions in previously unrecognized ways.

The azuki bean beetle *C. chinensis*, a major pest of stored pulses (Tuda, 2007), provides a useful model for studying multiple *Wolbachia* infections. Natural populations are predominantly coinfected with the CI-inducing strains wBruCon and wBruOri, although single infections also occur (Kondo et al., 2011, 2002). Elevated temperature can reduce fecundity, hatchability, and population size in this species (Tuda, 1993; Tuda & Shimada, 1993, 1995). Using the same infection lines and the same environmental treatments as in the present study, Leroy et al. (2025) recently showed that *Wolbachia* infection modified sex-specific parental investment under elevated temperature and CO_2_ (eT&CO_2_), anticipated future atmospheric conditions (IPCC, 2023). However, the effects of the same environmental conditions on the relative abundance of coinfecting strains, CI expression, and predicted infection persistence remain unknown. We therefore examined (i) how infection status, host sex, and eT&CO_2_ affect Con and Ori abundance; (ii) how infection status and eT&CO_2_ affect host life-history traits and CI expression; and (iii) how experimentally estimated fitness and incompatibility parameters, together with assumed maternal transmission values, influence predicted equilibrium infection frequencies.

## 2. Materials and methods

### 2.1. Insect lines

We used an isofemale line derived from the long-term laboratory strain jC of the azuki bean beetle *Callosobruchus chinensis* (Coleoptera: Chrysomelidae: Bruchinae), which has been maintained since 1937 (Utida, 1941) and is naturally coinfected with two *Wolbachia* strains, wBruCon (Con) and wBruOri (Ori). Con induces complete CI, whereas Ori induces partial CI (Kondo et al., 2002). From this isofemale line, we established three sublines: the naturally coinfected line (CO), a Con-only line (C) cured of Ori, and an uninfected line (U) cured of both strains. To establish the singly infected line, three pairs of adults were exposed for one generation to 800 µl of 0.05% tetracycline hydrochloride. Ori was consistently eliminated first, yielding the Con-only line; an Ori-only line could not be established. The uninfected line was subsequently produced from the Con-only line by treating three adult pairs for one generation with 800 µl of 0.25% tetracycline hydrochloride. Infection status was checked in offspring for two generations, and the cured lines were maintained for at least five generations before use in the experiments. The coinfected control line was exposed to the same volume of distilled water. Infection status was confirmed by PCR using the wsp forward primer wspF and the strain-specific reverse primers wspConR and wspOriR (Kondo et al., 2002). All lines were maintained at 30°C, 60 ± 8% relative humidity (RH), ∼500 ppm CO_2_, and a 16L:8D photoperiod on azuki bean, *Vigna angularis*. The long-term jC culture is maintained with a large census population (approximately 11,000 adults per generation), and previous work found little inbreeding depression in this culture (Tanaka, 1990).

### 2.2. Environmental treatments

Experiments were conducted in two environmental chambers. Control conditions were 30°C and ∼500 ppm CO_2_, whereas the combined elevated-temperature and elevated-CO_2_ treatment (eT&CO_2_) was 32°C and ∼1000 ppm CO_2_. Both chambers were maintained at 60 ± 8% RH with a 16L:8D photoperiod. Temperature, RH, and CO_2_ concentration were monitored daily with data loggers. Because temperature and CO_2_ were altered together, their individual effects cannot be separated in this experiment.

### 2.3. Quantification of *Wolbachia* relative abundance

Newly emerged pairs from the coinfected (CO) or Con-only (C) lines were allowed to oviposit on azuki beans in 5.5-cm-diameter Petri dishes under either control or eT&CO_2_ conditions until death. Four parental pairs were established for each infection-status × environment combination. Their offspring developed under the same environmental treatment. Before adult emergence, beans containing one or two larvae were isolated in 48-well plates. Virgin adults were collected within 24 h of emergence and stored at −60°C until DNA extraction.

DNA was extracted individually from 10 female and 10 male offspring from each coinfected or singly infected line reared under each environmental condition using a DNeasy tissue kit (Qiagen, Tokyo, Japan). The abundances of the *Wolbachia* strains were quantified using qPCR; the target gene *wsp* was amplified using strain-specific primers: TQwspC-F and TQwspC-R for Con and TQwspO-F and TQwspO-R for Ori (Ijichi et al., 2002). Primers specific to the host beetle nuclear gene *EF1α* were designed using Primer3 version 4.1.0 (Untergasser et al., 2012): qCcEF1α-F (5’-TGGCCACAGAGATTTCATCAAAA-3’) and qCcEF1α-R (5’-GTTCACACCAACAATGAGCTGT-3’). Negative controls containing water instead of the DNA template were included for each primer set. We used Luna Universal qPCR Master Mix (NEB) with the following cycling conditions: 95°C for 1 min, followed by 40 cycles of 95°C for 15 s and 60°C for 30 s. Amplification was analyzed with a StepOnePlus real-time PCR system (Applied Biosystems). Fluorescence readings were taken at the end of each cycle. The melting-curve protocol used a single cycle at 95°C for 15 s and at 60°C for 1 min. The temperature was increased from 60°C to 95°C at a ramp rate of 0.3°C s^−1^. The *C*_*T*_ values were normalized as 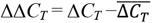, where Δ*C*_*T*_ is the difference between the *C*_*T*_ values for the target *Wolbachia* gene and the host gene and 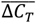 is the mean of Δ*C*_*T*_ for each *Wolbachia* strain in coinfected female hosts in the control environment. The abundance of *Wolbachia* DNA relative to host DNA was then calculated as 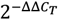 (Livak & Schmittgen, 2001).

qPCR efficiency was evaluated from standard curves generated using serial dilutions of samples with high target abundance. Three technical replicates were analyzed for each dilution. Estimated amplification efficiencies were 99.3% for the Con *wsp* primer set, 107.0% for the Ori *wsp* primer set, and 97.3% for the host *EF1α* primer set.

### 2.4. Host life-history traits

Newly emerged adults from the uninfected (U), Con-only (C), and coinfected (CO) lines were paired in all combinations of female and male infection status and allowed to oviposit on 5 g of azuki beans in 5.5-cm-diameter Petri dishes under either control or eT&CO_2_ conditions. Twenty parental pairs were prepared for each cross × environment combination. Fecundity, egg hatchability, development time, survival, and offspring sex ratio were recorded. One week after parental death, hatched and unhatched eggs were counted. Beans containing one or two hatched eggs were then isolated individually in 48-well plates for monitoring offspring emergence to estimate development time and survival (Fig. 1). Development time was recorded for the first three emerging females for each parental pair and means of the three females were calculated.

**Fig. 1.**
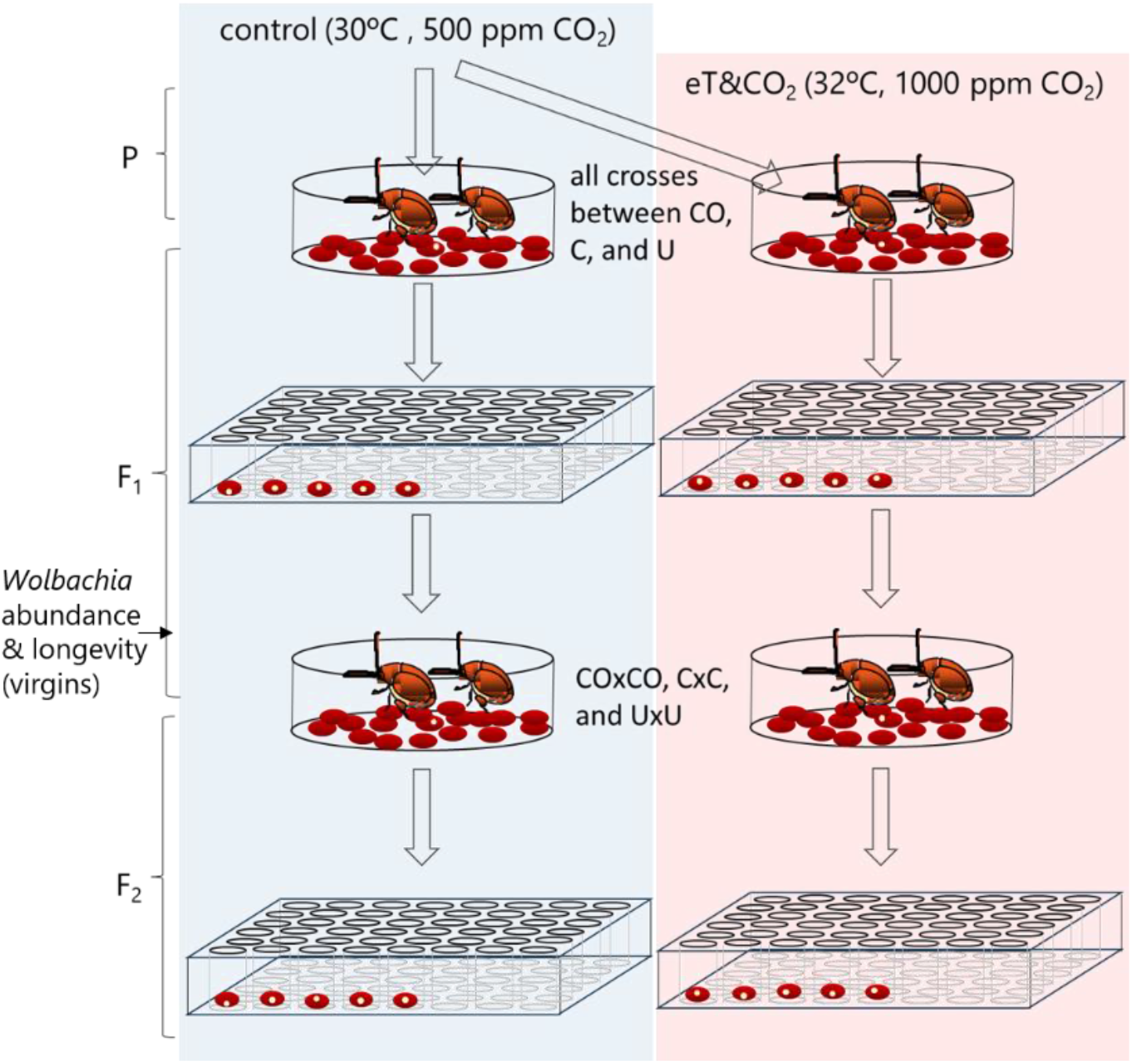
Experimental procedure. Coinfected (CO), Con-only (C), and uninfected (U) host (*Callosobruchus chinensis*) lines under control and combined elevated temperature and CO_2_ (eT&CO_2_) conditions.

Adult longevity was measured for offspring of the within-line crosses CO × CO, C × C, and U × U. Newly emerged parental pairs were allowed to oviposit for 48 h under either control or eT&CO_2_ conditions. Up to 20 beans bearing one hatched egg were sampled from each parental pair, isolated in 48-well plates, and maintained under the same environmental treatment (Fig. 1). Adult emergence and death were recorded daily. Eight parental pairs were used for each infection-status × environment combination. For the F2 assay, newly emerged F1 adults from CO × CO, C × C, and U × U crosses were paired within infection status and maintained under the same environmental treatment (Fig. 1).

### 2.5. Predicted equilibrium prevalence of CI-inducing *Wolbachia*

Equilibrium infection frequencies were predicted for Con (in singly infected or coinfected hosts) and Ori (in coinfected hosts) using the equations of Turelli (1994) and Jaenike (2009). These models include a higher stable equilibrium and a lower unstable equilibrium. Relative female fitness (*F*) was estimated from fecundity: CO × CO and C × C fecundity relative to U × U for the corresponding infection classes. Survival from incompatible crosses (*H*) was estimated as egg hatchability multiplied by larva–adult survival for U × CO (coinfection-associated incompatibility) or U × C (Con-associated incompatibility). Maternal transmission was not measured directly. We therefore treated *μ*, the fraction of uninfected eggs produced by infected females, as an assumed parameter and explored values from 0.1 to 0, corresponding to maternal transmission fidelities of 0.9–1.0. The resulting equilibria should therefore be interpreted as scenario predictions, rather than estimates, of infection frequencies. Sensitivity analyses were conducted by increasing and decreasing each parameter in turn (0.7 ≤ *F* ≤ 1.2, 0 ≤ *H* ≤ 0.2, or 0 ≤ *μ* ≤ 0.1).

### 2.6. Statistical analyses

Relative *Wolbachia* abundance was analyzed after log_10_ transformation. Con abundance and summed *Wolbachia* abundance (Con + Ori) were tested with general linear models including environment, host sex, infection status, and their interactions. Ori abundance in coinfected hosts was analyzed as a function of environment, host sex, log-transformed Con abundance, and their interactions. One extreme Ori-abundance observation from a coinfected female under control conditions was excluded from the analysis. Higher-order interactions with p > 0.15 were sequentially removed from the models.

Fecundity, development time, offspring number, proportion of females, and F2 egg–adult survival were analyzed using general linear models with parental infection status, environment, and their interaction as explanatory variables, after checking for the normality of model residuals. F1 egg–adult survival was analyzed by logistic regression. Adult longevity was analyzed using parametric survival models that included parental ID as an additional explanatory factor; Weibull and log-logistic distributions provided the best fits for females and males, respectively. Significant effects were followed by Tukey-type multiple-comparison tests. A significance level of *p* < 0.01 was used for all analyses. All analyses were performed in JMP version 19.1.0.

## 3. Results

### 3.1. *Wolbachia* abundance

Con relative abundance showed strong sex- and infection-status dependence, including significant environment × sex, environment × infection-status, sex × infection-status, and three-way interactions (Table 1). Under control conditions, Con abundance was exceptionally high in singly infected males but low in singly infected females. Under eT&CO_2_, Con abundance increased markedly in coinfected females, whereas it decreased in singly infected males relative to the control treatment (Fig. 2a). Across the other sex × infection-status combinations, Con abundance was intermediate. Thus, the response of Con to eT&CO_2_ depended on both host sex and whether Ori was present. In contrast, Ori abundance in coinfected hosts was lower under eT&CO_2_ than under control conditions (Table 1, Fig. 2a).

**Table 1.** General linear model results for the relative abundance of wBruCon (Con), wBruOri (Ori) in coinfected hosts, and summed Con + Ori abundance as functions of environmental treatment, host sex, infection status, and/or log-transformed abundance of the coinfecting strain. *p* values < 0.01 are shown in bold.

|  | Con |  |  | Ori in coinfected host |  |  | Summed Con and Ori |  |  |
|---|---|---|---|---|---|---|---|---|---|
|  | *df* | *F* | *p* | *df* | *F* | *p* | *df* | *F* | *p* |
| Environment | 1, 72 | 0.14 | 0.707 | 1, 35 | 50.82 | < **0.001** | 1, 72 | 2.06 | 0.155 |
| Sex | 1, 72 | 253.07 | < **0.001** | 1, 35 | 6.93 | 0.013 | 1, 72 | 243.22 | < **0.001** |
| Log(coinfecting strain abundance) | - | - | - | 1, 35 | 10.14 | **0.003** | - | - | - |
| Infection status | 1, 72 | 30.42 | < **0.001** | - | - | - | 1, 72 | 53.97 | < **0.001** |
| Environment × sex | 1, 72 | 26.21 | < **0.001** | - | - | - | 1, 72 | 24.56 | < **0.001** |
| Environment × log(coinfecting strain abundance) | - | - | - | - | - | - | - | - | - |
| Sex × log(coinfecting strain abundance) | - | - | - | - | - | - | - | - | - |
| Environment × sex × log(coinfecting strain abundance) | - | - | - | - | - | - | - | - | - |
| Environment × infection status | 1, 72 | 152.15 | < **0.001** | - | - | - | 1, 72 | 108.63 | < **0.001** |
| Sex × infection status | 1, 72 | 36.44 | < **0.001** | - | - | - | 1, 72 | 38.03 | < **0.001** |
| Environment × sex × infection status | 1, 72 | 30.21 | < **0.001** | - | - | - | 1, 72 | 31.05 | < **0.001** |

**Fig. 2.**
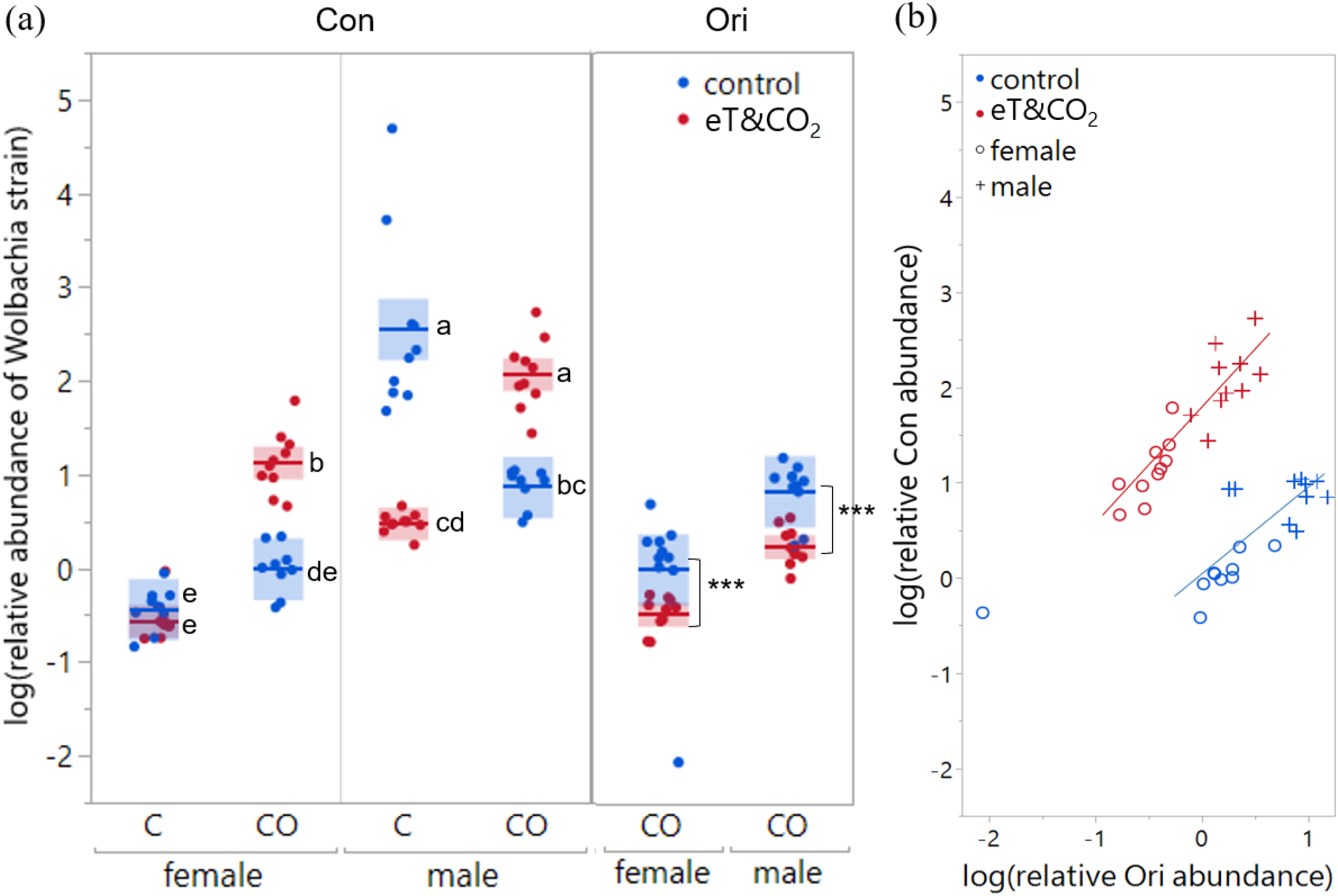
Relative abundance of *Wolbachia* strains under control and combined elevated-temperature and elevated-CO_2_ (eT&CO_2_) conditions. (a) Relative abundances of Con (wBruCon) and Ori (wBruOri) in coinfected (CO) and Con-only (C) beetles. Symbols show individual beetles; boxes show means with 95% confidence limits. Different letters indicate significant differences among Con groups; asterisks indicate environmental differences for Ori. (b) Relationship between Con and Ori relative abundance in coinfected beetles.

Within coinfected hosts, Ori abundance was positively associated with Con abundance [slope = 0.38 ± 0.12 (estimate ± SE) on the log–log scale; Table 1, Fig. 2b]. Summed *Wolbachia* abundance (Con + Ori) showed a pattern similar to that of Con, with significant effects of sex and infection status and significant two- and three-way interactions with environment (Table 1). Relative abundance was generally higher in males than in females and higher in coinfected than in singly infected hosts, except for the exceptionally high abundance observed in singly infected males under control conditions (Fig. 2a).

### 3.2. Host life-history traits

Parental fecundity differed among parental infection-status crosses (Table 2, Fig. 3a) but showed no overall effect of environment and no infection-status × environment interaction. Across crosses, fecundity tended to increase as the number of *Wolbachia* strains in the female decreased. The fecundity of coinfected females also varied with the infection status of the male, whereas the fecundity of Con-only and uninfected females showed little corresponding variation (Fig. 3a).

**Table 2.** Effects of parental cross infection status and environmental treatment (control or elevated temperature and CO_2_) on parental fecundity and F1 life-history traits. Parent ID was nested within infection status and environment for longevity analyses. L-R, likelihood-ratio test. *p* values < 0.01 are shown in bold.

|  | **Fecundity** |  |  | **Egg–adult survival** |  |  | **Female development time** |  |  | **Female offspring longevity** |  |  | **Male offspring longevity** |  |  |
|---|---|---|---|---|---|---|---|---|---|---|---|---|---|---|---|
|  | ***df*** | ***F*** | ***p*** | ***df*** | **L-R *χ*<sup>2</sup>** | ***p*** | ***df*** | ***F*** | ***p*** | ***df*** | **L-R *χ*<sup>2</sup>** | ***p*** | ***df*** | **L-R *χ*<sup>2</sup>** | ***p*** |
| Infection status | 8, 342 | 7.86 | < **0.001** | 8 | 217.88 | < **0.001** | 6, 256 | 54.94 | < **0.001** | 2 | 49.95 | < **0.001** | 2 | 74.43 | < **0.001** |
| Environment | 1, 342 | 2.15 | 0.14 | 1 | 6×10<sup>−7</sup> | 0.999 | 1, 256 | 110.50 | < **0.001** | 1 | 11.25 | < **0.001** | 1 | 86.35 | < **0.001** |
| Infection status × environment | 8, 342 | 1.61 | 0.12 | 8 | 0.19 | 1.000 | 6, 256 | 5.97 | < **0.001** | 2 | 4.71 | 0.10 | 2 | 16.68 | < **0.001** |
| Parent ID [infection status, environment] |  |  |  |  |  |  |  |  |  | 41 | 86.53 | < **0.001** | 41 | 76.49 | < **0.001** |

**Fig. 3.**
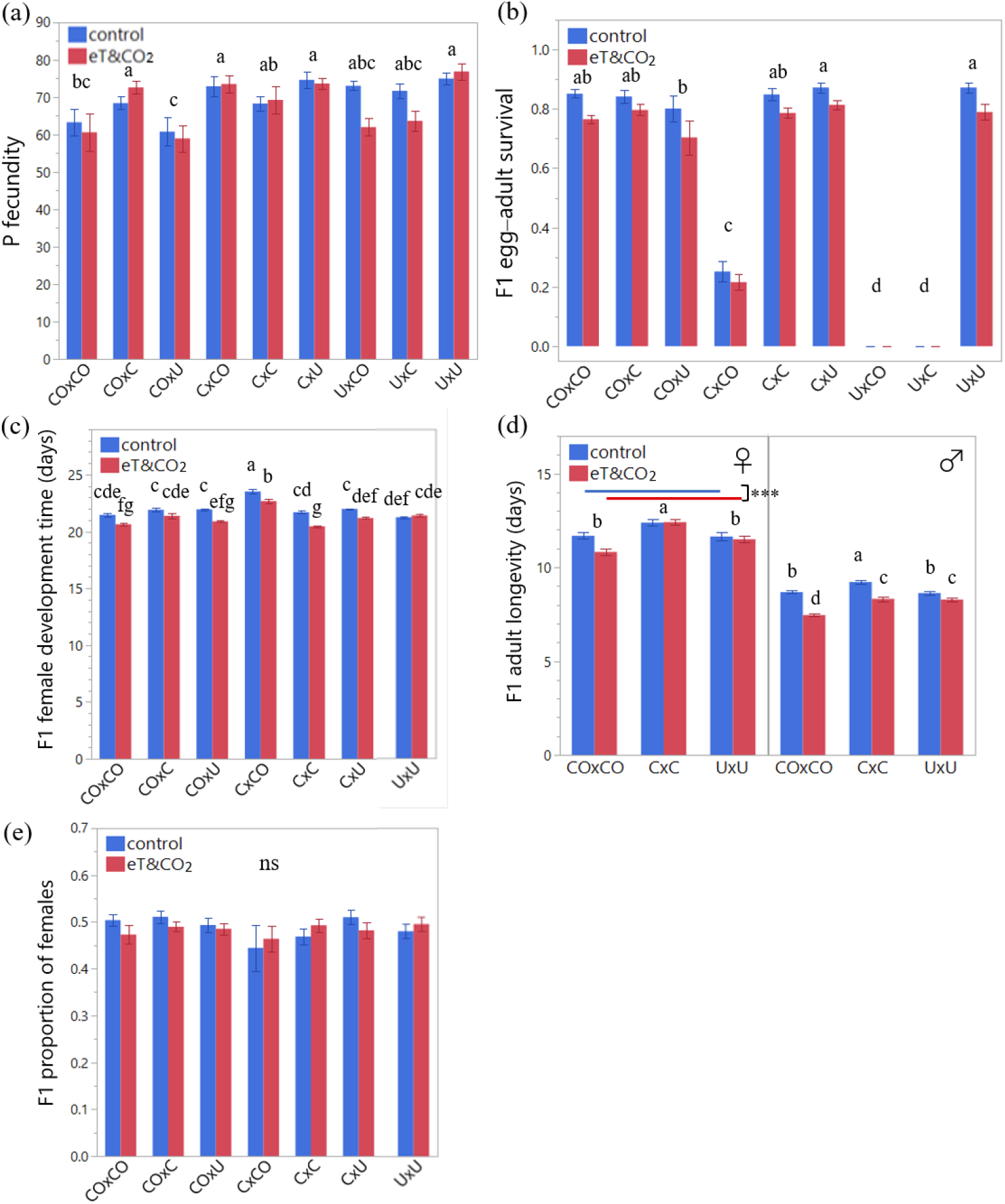
Parental fecundity and F1 life-history traits (mean ± SE) from female × male crosses among coinfected (CO), Con-only (C), and uninfected (U) host lines under control or combined elevated temperature and CO_2_ (eT&CO_2_) conditions. (a) Fecundity, (b) egg–adult survival, (c) female development time, (d) female and male adult longevity, and (e) proportion of females. Shared letters indicate no significant differences among relevant groups. ns, not significant; *** *p* < 0.001.

F1 egg–adult survival was affected by parental cross but not by environment and their interaction (Table 2, Fig. 3b). Exclusion of two incompatible crosses did not alter this qualitatively (infection status L-R *χ*^2^ = 52.95, df = 6, *p* < 0.001; environment L-R *χ*^2^ = 2.00, df = 1, *p* = 0.16; interaction L-R *χ*^2^ = 0.19, df = 6, *p* = 1.00). Survival was not different in most compatible crosses, except that CO × U survival was lower compared to C × U and U × U survival. As expected from Con-induced complete CI, the incompatible U × C and U × CO crosses produced no adult offspring and were excluded from subsequent analyses of traits requiring survivors. The C × CO cross showed reduced survival consistent with partial incompatibility associated with Ori.

F1 female development time was affected by parental cross, environment, and their interaction (Table 2, Fig. 3c). Development was generally faster under eT&CO_2_, although the magnitude of this response differed among crosses. Offspring from C × CO had comparatively long development times in both environments, whereas C × C offspring under eT&CO_2_ developed particularly rapidly.

Offspring sex ratio was not significantly affected by parental cross (*F*_6,258_ = 1.10, *p* = 0.36), environment (*F*_1,258_ = 0.15, *p* = 0.70), or their interaction (*F*_6,258_ = 0.67, *p* = 0.67); the proportion of females remained close to 0.5 across treatments (Fig. 3e).

F1 adult longevity was reduced under eT&CO_2_ in both sexes (Table 2, Fig. 3d). Parental infection status also affected longevity. Female offspring from C × C parents lived longer than those from CO × CO or U × U parents. For males, the parental infection-status × environment interaction was significant: under control conditions, males from C × C parents lived longer than those from CO × CO and U × U parents, whereas under eT&CO_2_ males from both C × C and U × U parents lived longer than those from CO × CO parents (Fig. 3d).

F1 fecundity was affected by the interaction between infection status and environment (Table 3, Fig. 4a). The clearest environmental decline occurred in the uninfected line, whereas fecundity of the coinfected and Con-only lines changed less between environments.

**Table 3.** Effects of infection status and environmental treatment (control or elevated temperature and CO_2_) on F1 fecundity and F2 life-history traits. *p* values < 0.01 are shown in bold.

|  | **Fecundity** |  |  | **Egg–adult survival** |  |  | **Female development time** |  |  |
|---|---|---|---|---|---|---|---|---|---|
|  | ***df*** | ***F*** | ***p*** | ***df*** | ***F*** | ***p*** | ***df*** | ***F*** | ***p*** |
| Infection status | 2, 114 | 3.34 | 0.039 | 2, 114 | 1.77 | 0.18 | 2, 112 | 0.59 | 0.56 |
| Environment | 1, 114 | 5.10 | 0.026 | 1, 114 | 21.30 | **< 0.001** | 1, 112 | 37.99 | **< 0.001** |
| Infection status × environment | 2, 114 | 6.56 | **0.002** | 2, 114 | 1.65 | 0.20 | 2, 112 | 1.88 | 0.16 |

**Fig. 4.**
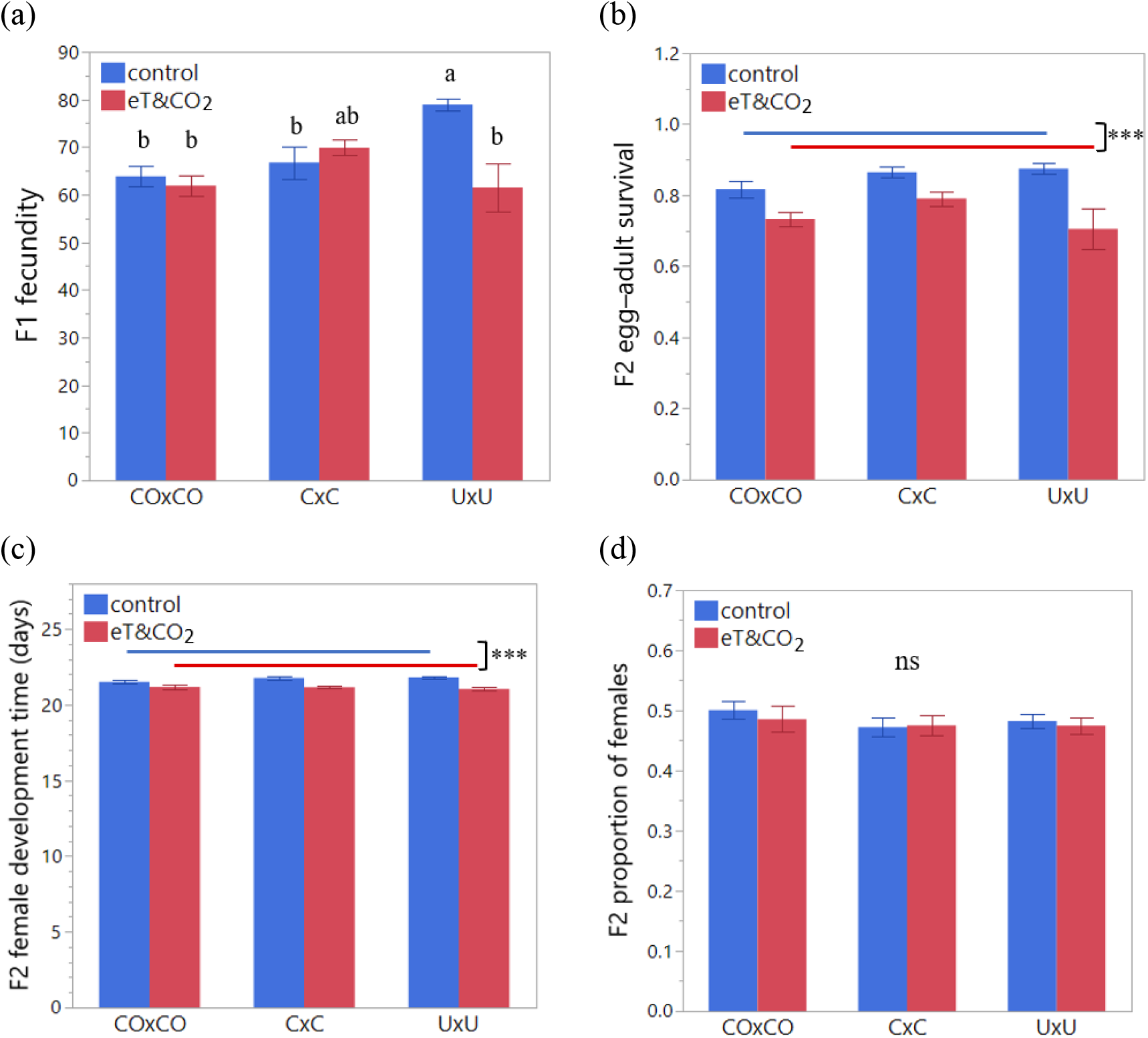
F1 fecundity and F2 life-history traits (mean ± SE) of within-line crosses under control or combined elevated temperature and CO_2_ (eT&CO_2_) conditions. (a) Fecundity, (b) egg–adult survival, (c) female development time, and (d) proportion of females. coinfected (CO), Con-only (C), and uninfected (U) host lines. Shared letters indicate no significant differences among relevant groups. ns, not significant; *** *p* < 0.001.

Egg–adult survival and female development time in the F2 generation were affected by environment, with lower survival and shorter development under eT&CO_2_ (Table 3, Fig. 4b,c). Offspring sex ratio was not significantly affected by parental line (*F*_2,112_ = 0.80, *p* = 0.45), environment (*F*_1,112_ = 0.27, *p* = 0.60), or their interaction (*F*_2,112_ = 0.16, *p* = 0.85); the proportion of females remained close to 0.5 across treatments (Fig. 4d).

### 3.3. Predicted equilibrium prevalence of the two CI-inducing *Wolbachia* strains

The parameter values used in the equilibrium models are summarized in Table 4. Using these parameter values, the equilibrium model predicted stable equilibria staying at 1.0 for both the coinfected and Con-only infection classes across environments. By contrast, under eT&CO_2_, the unstable equilibrium was higher for the coinfected class using the first-generation estimates (P’s *F* and F1 *H*) and assumed values (P–F1 *μ*) (Fig. 5a), but lower for both infection classes using second-generation estimates (F1 *F* and F2 *H*) and assumed values (F1–F2 *μ*) (Fig. 5b). Lower relative fecundity (parental *F)* in the first generation and higher F1 *F* in the second generation contributed to these shifts in unstable equilibria. Consequently, the interval between the stable and unstable equilibria narrowed under eT&CO_2_ using first-generation estimates (Fig. 5a) but widened using second-generation estimates (Fig. 5b). Within the ranges examined, sensitivity analyses indicated that changes in *F* had a greater effect on the unstable equilibrium than changes in *H* or *μ*. Therefore, the higher *F* value is predicted to improve *Wolbachia* persistence in the second generation.

**Table 4.** Parameter values used to predict equilibrium frequencies of the CI-inducing *Wolbachia* strains Con and Ori using the equations of Turelli (1994). *F*, relative fitness of infected females; mother *F*, parental-generation *F* estimate; daughter *F*, F1-generation *F* estimate; *H*, survival of offspring from incompatible crosses (CI strength = 1 − *H*); *μ*, fraction of uninfected eggs produced by infected females (1 − transmission fidelity). Maternal transmission was not measured; *μ* was therefore explored over the assumed range. eT&CO_2_, combined elevated temperature and CO_2_ treatment.

|  | **control** |  |  |  | **eT&CO<sub>2</sub>** |  |  |  |
|---|---|---|---|---|---|---|---|---|
|  | **Mother *F*** | **Daughter *F*** | ***H*** | **Assumed *μ* range** | **Mother *F*** | **Daughter *F*** | ***H*** | **Assumed *μ* range** |
| Con + Ori | 0.844 | 0.809 | 0 | 0–0.1 | 0.789 | 1.006 | 0 | 0–0.1 |
| Con | 0.911 | 0.845 | 0 | 0–0.1 | 0.901 | 1.135 | 0 | 0–0.1 |

**Fig. 5.**
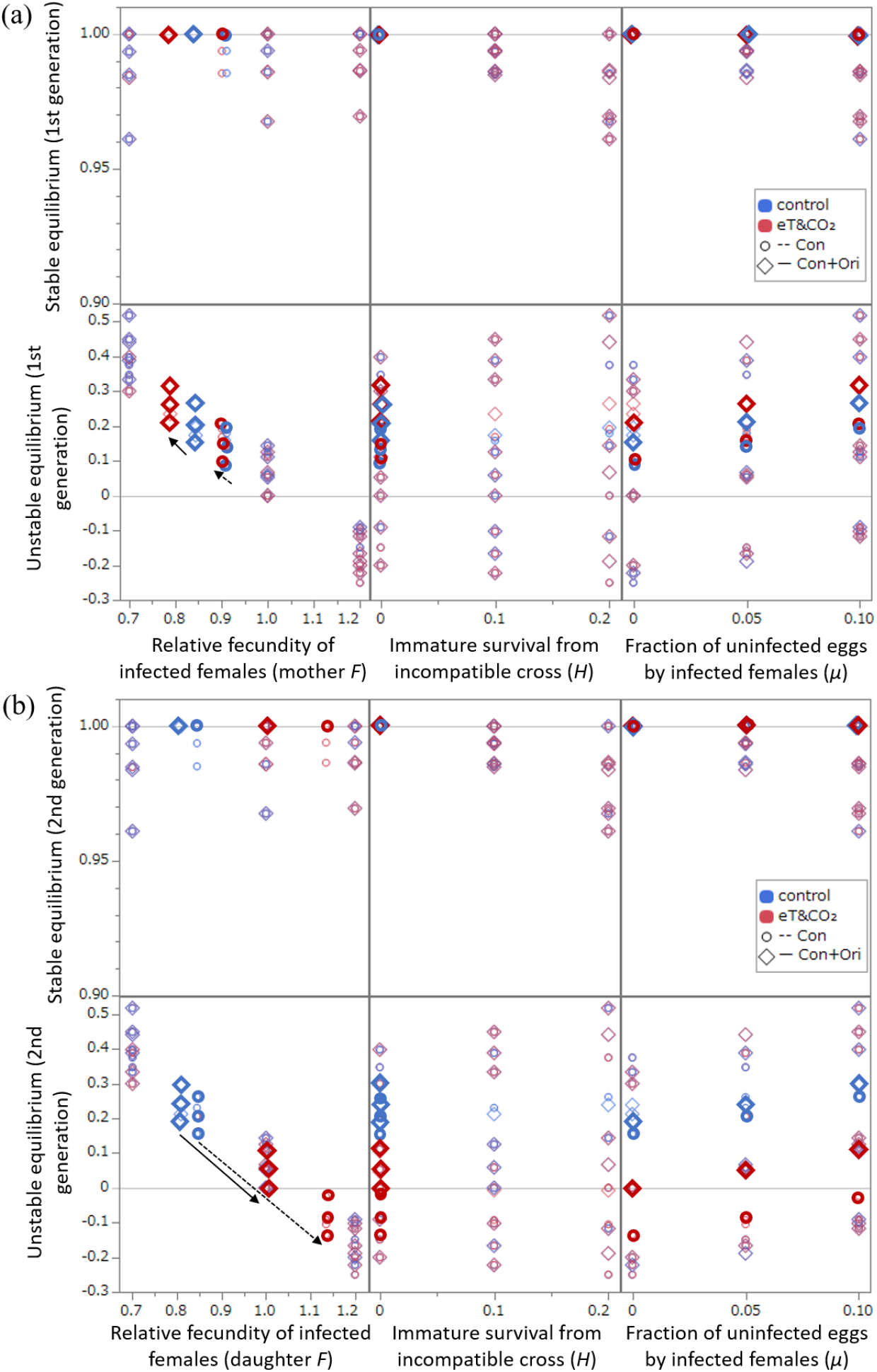
Predicted equilibrium frequencies of the two CI-inducing *Wolbachia* strains based on *F* (relative fitness of infected females), *H* (survival from incompatible crosses; CI strength = 1 − *H*), and μ (fraction of uninfected eggs produced by infected females) for (a) first generation and (b) second generation. *F* and *H* are observed values. *μ* is varied within an assumed range. Thick symbols indicate the estimated parameter values for *F* and *H* and assumed values for *μ* (Table 4); thinner symbols show sensitivity analyses. Arrows indicate the contribution of *F* to unstable equilibrium changes from control to combined elevated temperature and CO_2_ (eT&CO_2_) conditions.

## 4. Discussion

The principal result of this study is that the two coinfecting *Wolbachia* strains responded differently to the same environmental perturbation. Under eT&CO_2_, Ori relative abundance declined whereas Con abundance increased in coinfected females and decreased in singly infected males. Con abundance therefore depended jointly on host sex, environmental treatment, and the presence of Ori. In contrast, environmental effects on host life-history traits were trait dependent and those on some traits, such as F1 female development time, male longevity, and fecundity, were different among infection statuses. Together, these findings indicate that environmental change can have pronounced effects on the internal dynamics of coinfecting symbionts without producing similarly large effects on host performance.

The equilibrium analysis illustrates how environmentally induced changes in relative fitness could influence the theoretical stability of infected populations. Under eT&CO_2_, the unstable equilibrium increased for the coinfected class when first-generation estimates were used but decreased for both infection classes when second-generation estimates were used. Consequently, the interval between stable and unstable equilibria narrowed in the first generation but widened in the second. In the Turelli (1994) framework, a wider interval indicates greater resistance to perturbations that reduce infection frequency because a larger decline is required before the infection falls below the unstable threshold. Thus, the increased relative fitness of F1 infected females may improve *Wolbachia* persistence in the second generation under environmental stress. Such a mechanism could contribute to the persistence of high coinfection frequencies reported in natural populations, although field transmission and fitness parameters remain to be quantified (Kondo et al., 2011). However, because maternal transmission was not measured directly, these equilibrium values should be interpreted as scenario predictions rather than estimates of field prevalence.

The positive association between Con and Ori abundance in coinfect ed beetles is consistent with shared host-level regulation or similar responses to host physiological conditions. At the same time, the contrasting environmental responses of the two strains indicate that coinfecting *Wolbachia* need not respond uniformly to environmental change. Con and Ori differ in within-host distribution (Ijichi et al., 2002), which may facilitate coexistence while permitting strain-specific responses. The exceptionally high Con abundance in singly infected males under control conditions further suggests that Ori may constrain Con proliferation in a sex-dependent manner. Similar sex- and tissue-specific variation in coinfecting symbiont dynamics has been reported in *D. simulans* (Jones et al., 2022). Although the present data do not distinguish direct strain interactions from host-mediated regulation, they demonstrate that environmental change can alter the relative abundance of coinfecting strains and thereby potentially influence infection stability and symbiont-associated phenotypes.

Despite substantial shifts in relative abundance, incompatibility patterns remained stable across environments. Complete incompatibility associated with Con and partial incompatibility associated with Ori were both maintained under eT&CO_2_. This result is consistent with previous evidence that CI expression can remain robust across a range of *Wolbachia* densities (Gharabigloozare & Bleidorn, 2022). The reduction in hatchability observed in C × CO crosses was greater than that reported by Kondo et al. (2002), possibly reflecting differences in host genetic background or experimental conditions.

Host responses to eT&CO_2_ were trait dependent and varied across generations. Development was accelerated and adult longevity was reduced under eT&CO_2_, whereas effects on survival were most evident in the F2 generation. Fecundity of the parental generation was comparatively insensitive to the environmental treatment, potentially because newly emerged *C. chinensis* females already possess a large complement of mature oocytes. Infection status also influenced several life-history traits and modified environmental responses for some of them. Most interestingly, while infected lines retained their F1 fecundity, F1 fecundity in the uninfected line sharply declined under eT&CO_2_, suggesting that infected lines maintained relative fitness more effectively.

These patterns complement the parental-investment response reported by Leroy et al. (2025), who found that coinfected parents increased male egg size under the same environmental treatment. Taken together, the two studies suggest that *Wolbachia*-associated responses to environmental change may be expressed more strongly through parental effects and relative fitness differences than through large changes in mean fecundity alone. Coinfected parents generally produced fewer eggs than uninfected parents, while the Con-only line was often intermediate, consistent with a reproductive cost of infection in this host background. Likewise, the shorter lifespan of coinfected offspring relative to Con-only offspring may indicate an additional cost associated with Ori or coinfection.

In conclusion, elevated temperature and CO_2_ altered the abundance of coinfecting *Wolbachia* strains and modified their predicted infection-frequency thresholds in a host population. Because infected lines retained relative fitness advantages under eT&CO_2_ while CI remained effective, environmental change may influence both the dynamics of natural *Wolbachia* infections and symbiont-induced resistance against pesticides (Miksanek & Tuda, 2023). These findings highlight the need to incorporate environmental variation when predicting the ecological consequences of host–symbiont associations.

## CRediT authorship contribution statement

Phy Phyu San: Conceptualization, Formal analysis, Writing – original draft, Visualization, Methodology, Investigation. Ryosuke Fujita: Methodology, Formal analysis. Julia Ferrari: Writing – review & editing. Midori Tuda: Conceptualization, Methodology, Formal analysis, Writing – original draft, Writing – review & editing, Visualization, Supervision.

## Funding

MT was supported by KAKENHI (19K06840) from JSPS (Japan Society for the Promotion of Science) and by the Initiative for Realizing Diversity in the Research Environment from MEXT (Ministry of Education, Science and Technology) to Kyushu University.

## Declaration of competing interests

The authors declare no known conflicts of interest or competing interests.

## Acknowledgments

We thank Masato Hino for his suggestion of the literature and James Miksanek and Jamie Alves for their comments on our manuscript.

## Data availability

The datasets associated with the current study are available from the primary corresponding author on reasonable request.

## Notes

### Competing Interest Statement

The authors have declared no competing interest.

